# Regular *Plasmodium falciparum* importation onto Bioko Island, Equatorial Guinea, hampers malaria elimination from the island

**DOI:** 10.1101/2024.12.19.629489

**Authors:** Thomas C. Stabler, Ankit Dwivedi, Bing Guo, Biraj Shrestha, Sudhaunshu Joshi, Matilde Riloha Rivas, Olivier Tresor Donfack, Carlos A. Guerra, Guillermo A. García, Claudia Daubenberger, Joana C. Silva

## Abstract

The Bioko Island Malaria Elimination Project (BIMEP) has made significant progress in reducing the prevalence of *Plasmodium falciparum* on Bioko Island, Equatorial Guinea. However, like other malaria endemic islands like São Tomé and Principe and Zanzibar, Tanzania, elimination efforts are hampered by imported infections. In an effort to understand the local transmission dynamics and the influence of importation on Bioko Island’s *P. falciparum* population, whole-genome sequences were generated from field samples collected during the BIMEP’s 2019 Malaria Indicator Survey (MIS). Within the sub-Saharan African context, we observed Bioko Island parasites did not significantly differentiate from nearby continental neighbors. Among Bioko infections, within-host diversity and the quantity of polyclonal infections appear similar to an area of moderate malaria transmission. However, we observed higher than expected genetic diversity among Bioko parasites, similar to high transmission areas, suggesting imported strains are contributing to transmission on the island. Among Bioko’s closest geographical neighbors, the flow of parasites with Bioko appeared more pronounced with the Gabonese parasite population, implying more importation may be coming from this region than others. Overall, despite significant investment in malaria control, results illustrate the challenges of eliminating malaria without both interrupting local transmission and accounting for importation from higher transmission areas, likely due to human migration. For there to be sustained progress towards elimination, the BIMEP needs, if feasible, to conduct targeted interventions of outgoing/incoming travelers, and ideally expand malaria control interventions to the continental region of Equatorial Guinea.

**Importance:** *Plasmodium falciparum* accounts for the majority of malaria deaths globally, with over 500,000 estimated deaths in 2023, predominantly in sub-Saharan Africa, despite strong investment in control and elimination interventions. Incorporating sequencing technologies into malaria surveillance is viewed as a powerful tool to improve control strategies, including identifying parasite transmission pathways. Here, we provide the first genomic characterization of *P. falciparum* on Bioko Island and Equatorial Guinea since malaria control began in 2004, and use these data to better understand the contribution of neighboring regions to Bioko’s *P. falciparum* population. Results highlight the need to account for off-island contributors to transmission and to understand how unmitigated transmission in neighboring regions can hamper progress. This study furthers our understanding of how the flow of parasites between regions impacts infectious disease control and provides foundational genomic data in a previously undescribed region that can be used to inform malaria elimination efforts.

## Background

In the effort to eliminate malaria, islands hold a distinct advantage over malaria-endemic areas with land borders: their geographic isolation should, in theory, provide a barrier to parasite importation. With the introduction of effective control/prevention methods, this should facilitate rapid progress to malaria elimination. This has been observed in Sri Lanka and Cabo Verde, where no autochthonous cases have been reported since 2009, in the former, and 2017, in the latter, and which were declared malaria free in 2016 and 2024, respectively (1-3). Unfortunately, for other island contexts, like Zanzibar and São Tomé and Principe, malaria continues to persist despite significant decreases in malaria prevalence (4, 5). Parasite migration, through host and/or vector movement, poses a significant obstacle to malaria elimination (6, 7). Critically, the identification of factors preventing malaria elimination on island settings could provide valuable evidence for malaria control programs to distribute their resources.

Bioko Island, Equatorial Guinea, is located 32 km west of the coast of Cameroon and has historically high levels of malaria transmission (8) (**Figure 1A**). Intensive malaria control interventions have been conducted on the island since 2004 by the Bioko Island Malaria Elimination Project (BIMEP) (formerly the Bioko Island Malaria Control Project – BIMCP), which reduced malaria prevalence from 43.3% in the early 2000s to 10.5% by 2016 (8). Despite continued interventions, which also resulted in significant reductions in malaria mortality on the island (8, 9), malaria prevalence has fluctuated around 10-12% since 2016 (9, 10). Epidemiological, questionnaire-based studies by BIMEP demonstrate strong associations between on-island infections and recent travel to mainland Equatorial Guinea, suggestive of case importation, and identifying host movement as a potential source of parasite immigration to the island (9, 11-13). Further, as part of the COVID-19 pandemic response, a travel moratorium was implemented country-wide in Equatorial Guinea and without which it is estimated malaria infectious would be 9% higher in communities with more reported travel (14). However, the contribution of parasite migration to the composition of the Bioko Island parasite population remains to be validated and characterized molecularly.

**Figure 1.**
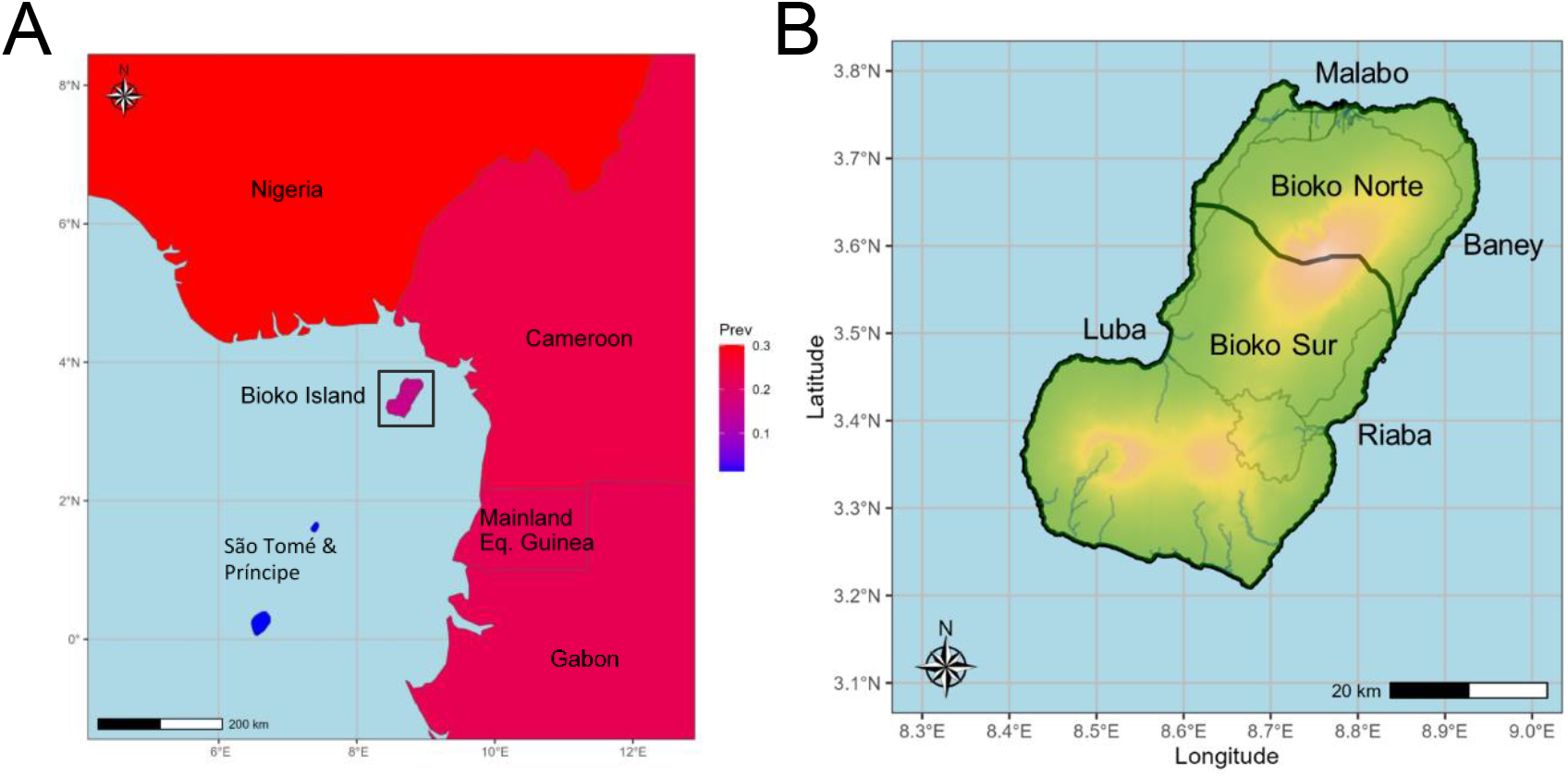
Map of Bioko Island, Equatorial Guinea. A. Geographical location of Bioko Island within the Gulf of Guinea. Color gradient denotes the estimated country-wide *P. falciparum* prevalence in 2019 reported in the 2023 World Malaria Report. Bioko Island color was amended to show island-wide prevalence as measured from the 2019 MIS. B. Detail of Bioko Island with province borders (black lines), roads (grey lines), rivers (blue lines) and elevation (green/red gradient – red denotes higher elevation).

The mainland region of the country, Rio Muni, has higher malaria prevalence than Bioko Island and has not had significant malaria control campaigns since 2011 due to funding constraints (15). Cameroon and Gabon, neighbors flanking Rio Muni, are hyper-endemic regions with year-round transmission and high parasite genetic diversity (1, 16, 17). If migration from the continent to Bioko Island is high, the island’s *Plasmodium falciparum* population, the predominant parasite species causing malaria (1), is expected to have a similarly high genetic diversity and little differentiation from Cameroon and Gabon. However, if the ocean forms a significant geographical barrier, and importation does not significantly contribute to Bioko’s malaria transmission and genetic diversity, then its *P. falciparum* population might differ significantly from those on the mainland.

Bioko Island is composed of two provinces (Bioko Norte and Bioko Sur) and four districts (Malabo, Baney, Luba and Riaba) (**Figure 1B**); Malabo City, the capital of Equatorial Guinea, located in Malabo district, Bioko Norte, harbors 90% of the island’s population of 270,000 people (9, 18, 19). Historically, the greatest reductions in malaria burden have been achieved in rural communities, especially in the Baney and Luba districts (8). However, rural residents also account for most of the on-island travel, Malabo being the primary destination for employment and education (12). Considering the disparate human distribution and reported travel, more than one parasite subpopulation may exist. Conversely, importation and inter-island connectedness may be sufficiently high as to prevent differentiation and the formation of subpopulations.

The use of whole genome sequence data to estimate population genetic diversity and relatedness among *P. falciparum* strains can provide a highly nuanced assessment of the impact of control interventions over time and illuminate parasite transmission routes (20, 21). Consequently, WHO encourages malaria control programs to incorporate parasite genome sequencing technologies among their molecular surveillance techniques (22). Previous studies have investigated demography of *P. falciparum* populations using whole genome data in sub-Saharan African (23); however, the vastness of this region has resulted in geographic “pockets” with incomplete characterization of circulating genetic diversity. Especially in areas with active malaria control campaigns but limited history of molecular surveillance, characterization of *P. falciparum* genome-wide genetic variation and estimating genetic relationships among isolates can provide insights into the response of a parasite population to ongoing interventions and provide valuable data to inform future approaches to parasite control (20, 24-26).

Here, we aim to characterize the genetic diversity and population structure of *P. falciparum* on Bioko Island and determine its relationship with parasite populations in neighboring regions on the mainland. Dried blood spot (DBS) samples obtained from a subset of participants in the BIMEP’s 2019 malaria indicator survey (MIS) were used to generate whole-genome sequence (WGS) data. Differentiation between Bioko Island and continental *P. falciparum* parasite populations was estimated, local transmission dynamics studied, by describing island-specific markers and relatedness between *P. falciparum* isolates.

## METHODS

### Sample collection

DBS samples were collected on Whatman filter papers (GE healthcare Ltd, Forest farm, Cardiff, UK; Product code: 11962089) from August to October 2019. On each filter paper are blood spots of 0.5 inches in diameter (n=4), each spot representing a volume of approximately 50 μl of blood. Survey and laboratory data used in this study include: gender, rapid diagnostic test (RDT) diagnosis (*P. falciparum* or mixed infection), qPCR diagnosis (*P. falciparum* or mixed infection), travel history (previous two months), location (urban or rural, or Malabo, Baney, Luba, Riaba districts), age group in years (<5, 5-15, 15-18, >18), and parasite density by qPCR (Low: Cq ≥ 20: High: Cq < 20). Household sampling was based on primary sampling units (PSUs) constructed from 1×1 km map-areas that make up the Bioko mapping grid (27). PSUs were assigned to either a rural or an urban stratum based the density of households within a community, with 25% and 5% of households sampled from each, respectively. All MIS participants (or their legal guardians, for participants <18 years of age) provided informed consent. Filter papers were selected for DNA extraction from 202 individuals with a positive RDT and reported fever.

Additionally, publicly available WGS data generated by the MalariaGEN *Plasmodium falciparum* Community Project (28) were downloaded from the Sequence Read Archive (SRA). Samples were selected from West, Central and East African countries as representative of their respective continental regions (**Supplementary File S2**).

### DNA extraction

A DNA extraction method based on guanidine and silica purification protocols, and developed by the Malaria Research Program, at the University of Maryland Baltimore (UMB), was applied to selected DBS filter paper samples as described before (29). Briefly, one circle/DBS was cutout, incubated, and submerged in lysis buffer for 2 hours at 65°C. Samples then underwent two washes before extracting DNA with TE buffer. Extracted DNA material was stored at -80°C until use.

### Polymerase chain reaction (PCR)

The Qiagen QuantiTect® Multiplex PCR was used to conduct qPCR (Qiagen Sciences, Germantown, Maryland, USA). The master mix was prepared according to manufacturer’s instructions, but adapted to exclude ROX or UDG. RNAseq free water was used in all PCR reactions. The following PCR program was used: [1] 20 minutes at 50°C; [2] 15 minutes at 95°C; [3] 45 seconds at 94°C; [4] 75 seconds at 60°C. Steps 3 and 4 were repeated 45 times. For each sample, all PCR reactions were performed in duplicate. Samples were considered positive if the mean quantification cycles (Cq) of duplicate qPCR reactions was Cq < 40. If one sample result was reported as RDT positive and the corresponding PCR result was negative, the assay was repeated to make a definitive conclusion, if necessary.

### Selective Whole Genome Amplification (sWGA)

Selective whole genome amplification (sWGA) was applied on extracted DNA with highest parasite density by Cq, as previously described (30, 31). In short, selected samples underwent a vacuum filtration step and were then amplified in 0.2 mL 96-well PCR plate with the following reaction mixture: 1X BSA, 1mM dNTPs, 2.5 μM of amplification primers, 1X Phi29 reaction buffer and 20 units of Phi29 polymerase. Primers are the same used in Oyola SO, et al. that preferentially bind at adequate distance and amplify *P. falciparum* genomes (31). From each extraction, 17 μL of the DNA sample was added to the reaction mixture for a total of 50 μL final volume. Amplification occurred in a thermocycler with the following stepdown protocol: 35°C for 20 minutes, 34°C for 10 minutes, 33°C for 15 minutes, 32°C for 20 minutes, 31°C for 30 minutes, 30°C for 16 hours. The stepdown protocol was followed with a heating step at 65°C for 20 minutes and cooled to 4°C.

### Whole genome sequencing

Preparation of genomic DNA libraries was previously described (32). To summarize, genomic DNA libraries were generated from amplified samples using the KAPA Library Preparation Kit (Kapa Biosystems, Woburn, MA), and then fragmented to approximately 200 base pair (bp) lengths. A modified version of the manufacturer’s protocol was used. AMPure XT beads were utilized to inform the library size selection. DNA concentration and fragment size was conducted using the DNA High Sensitivity Assay on the LabChip GX (Perkin Elmer, Waltham, MA) tool. All sample libraries were uniquely barcoded, pooled and sequenced on 150 bp paired-end Illumina NovaSeq 6000 run (Illumina, San Diego, CA).

### Read mapping and identification of single nucleotide polymorphisms (SNPs)

Raw fastq files were mapped to the reference *P. falciparum* genome, 3D7, using bowtie2 (33). BAM file processing followed GATK Best Practices (34, 35). Coverage and depth estimates from reads were generated using bedtools (36). Variant calling of each sample was conducted utilizing the Haplotype Caller toolkit to generate genomic variant call format files (GVCF) and perform joint SNP (single nucleotide polymorphisms) calling. When appropriate, diploid calls were allowed since polyclonal infections were expected; otherwise major alleles were called (70% threshold) to genotype the most prevalent strain in the infection (polymorphic sites that did not reach the threshold were set to missing). Variant calls were filtered to omit potential false positive results with the following criteria: DP<5, FS>14.5, MQ<20.0, QUAL<50. Additional filtering was performed to exclude rare allele events (frequency less than 0.05%), sites with >10% missingness, samples with >20% missing genotype values (37).

### Genetic Diversity

Genetic variation in each geographic region was estimated by nucleotide diversity (*π*) (Vcftools v.0.1.16), the average pairwise difference between samples. Within-host diversity was measured by the F-statistic *F*_WS_ (38) using the R package moimix (v0.0.2.9001, https://bahlolab.github.io/moimix/), where diversity of *P. falciparum* in each sample was compared against the diversity of the entire sample set. Infections were considered polyclonal if *F*_WS_ < 0.95. *P*-values were calculated using Chi-squared test to determine statistical significance for differences in proportions of polyclonal infections, as determined by *F*_WS_, between African countries.

### Population structure

Principal components analysis (PCA) was applied to a data set of biallelic SNPs passing filtering criteria, to investigate the extent to which geographic origin contributes to differences between isolates at the genome-wide level. Clustering by PCA was estimated using the R package SNPRelate (v.1.28.0; https://github.com/zhengxwen/SNPRelate) (39, 40). Admixture analysis (ADMIXTURE v1.2) was used to obtain an estimate of contributions of inferred ancestral subpopulations to each sample. PCA and admixture data sets were pruned for sites in linkage disequilibrium (LD) (window size of 5 kbp, *r*^2^≥0.2) prior to analysis. Among samples with a clonal pair (i.e. nearly identical genomes), one sample was selected to represent the clonal group, and the other(s) excluded from the analysis. Chi-squared test was used to measure differences in composition among each inferred ancestral subpopulation between countries.

### Genetic differentiation

Wright’s fixation index, *F*_ST_, was applied to measure overall differentiation (mean *F*_ST_) between sampled parasite populations from different geographic regions, and to identify SNPs contributing to differences between populations (41). Statistical significance of *F*_ST_ values were estimated empirically, with 5,000 permutations of samples by geographic region, using custom scripts.

### Relatedness

Overall relatedness between strains, as measured by identity-by-descent (IBD), was estimated using hmmIBD (version 2.0.0; https://github.com/glipsnort/hmmIBD) (42). Prior to conducting IBD, samples were deconvoluted using dEploid v0.6-beta (43). The strain accounting for the highest proportion of the sample was selected for downstream analysis. Previously published Python-based scripts were used to process files and run hmmIBD, and infomap (44), a community detection software, excluding IBD segments under 2cM and highly conserved IBD peaks to minimize bias (45). Samples were considered related (siblings or clonal) if >25% of genomes were IBD (46). Wilcoxon rank sum test was used to measure the difference in IBD distribution between sample sets.

### Visualization

The R package sf (47) were used to generate maps of Bioko. R (v4.1.3) was used with all appropriate R-based packages, such as ggplot2, to generate figures.

## RESULTS

### Sample characteristics

The DNA Cq of the original 160 Bioko Island samples analyzed ranged between 24.3 and 38.9 (mean Cq = 33.7). Samples with highest parasite density were prioritized for sWGA (range: 24.3 – 38.2; mean Cq = 32.9). Whole genome shotgun sequencing (WGS) data was successfully generated for 90 samples. Of these, fifty-two samples were collected in urban communities compared to 38 from Bioko’s rural areas (**Figure 2**). An average of 28,276,493 total reads were generated per sample; on average, across samples, 75.4% of the reference 3D7 genome was genotyped with at least 5X read coverage (Supplementary Figure S1). Of the 90 samples, 16 samples were excluded due to inadequate quantity of mapped reads (< 8 million) and/or high missingness (>20% SNPs missing).

**Figure 2.**
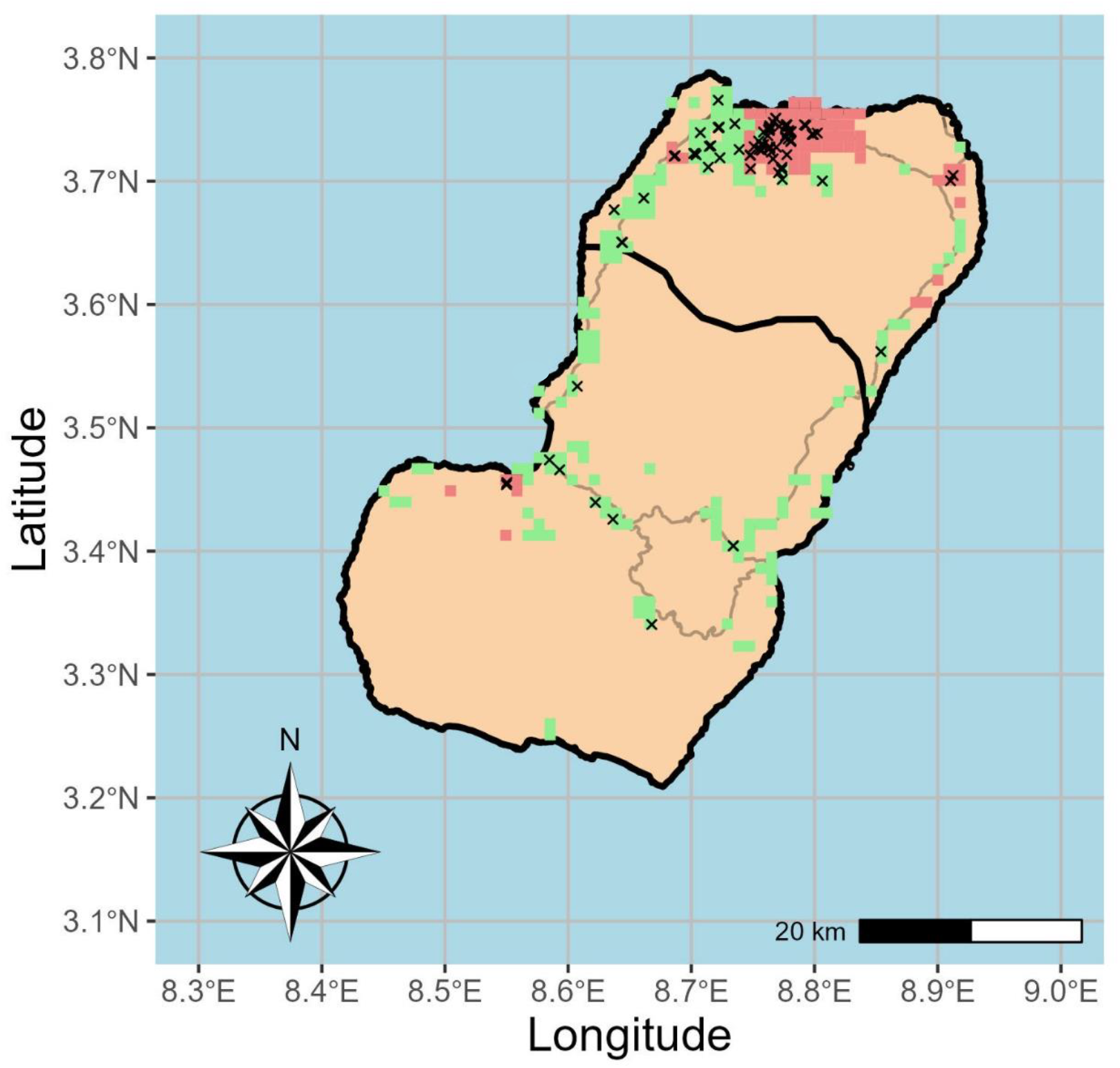
Urban and Rural Communities on Bioko Island. Map of Bioko Island with communities color-marked as urban (red) or rural (green). The urban/rural status was determined by the density of households within a community (12). Black crosses represent geographical location of collection of all sequenced samples (n=90).

Bioko Island WGS data sets were combined with 992 sub-Saharan African WGS data sets after excluding samples based on genotype missingness (n=25), and highly related, or clonal, clusters, where only one sample was included to represent the clonal group (n=14). Public samples were selected from African countries representative of the continent’s West (Guinea, Mali, Burkina Faso, and Nigeria), Central region (Cameroon, Gabon and Democratic Republic of Congo) and East (Kenya, Tanzania, Malawi, and Mozambique). Variable sites in the highly polymorphic sub-telomeric genomic regions were excluded due to reduced accuracy (48, 49). After calling major alleles, a total of 1,076,963 biallelic SNPs were included, of which 375,921 (34.9%) were intergenic, 205,240 (19.1%) were synonymous and 426,551 (39.6%) were non-synonymous. Overall, average read coverage across all sites was 63.0X (59.9X for Bioko Island samples).

### Bioko island P. falciparum population harbors high levels of genetic variation

Island settings with strong malaria elimination policies have seen a decrease in genetic diversity, and accompanying expansion of clonal parasites (50-52). To describe the genetic variation of the *P. falciparum* population among Bioko Island samples (n=74), nucleotide diversity and within-host diversity (*F*_WS_) were measured. Nucleotide diversity per site, estimated among variable sites only, was similar between Bioko district subgroups (*π*_Malabo city_ = 0.0088 ± 0.04; *π*_Malabo suburbs_ = 0.0086 ± 0.05; *π*_Baney_ = 0.0076 ± 0.06; *π*_Luba_ = 0.0079 ± 0.05). Nucleotide diversity was high among Bioko samples compared with that observed in nearby countries (all sites = 0.0087; nonsynonymous sites = 0.0088; synonymous sites = 0.0085) (Supplementary Figure S2). Within-host diversity measured by *F*_WS_ revealed the majority (46 out of 74) of isolates in Bioko Island were polyclonal (mean *F*_WS_ = 0.86) (**Figure 3**). While most polyclonal infections (n=41) originated in Malabo communities, there was no significant difference in distribution of polyclonal infections between Malabo, Baney and Luba communities (*p*-value = 0.31). Further, the proportion of complex infections in Bioko Island did not significantly differ from the proportions seen in Cameroon (*p*-value = 1) or Gabon (*p*-value = 0.56), the closest African countries on the African mainland. When analyzing *F*_WS_ results by epidemiological subgroup, the odds of carrying a polyclonal infections was higher for children and travelers, consistent with previous epidemiological associations (9, 12, 19). However, this trend was not statistically significant (Supplementary Table S1).

**Figure 3.**
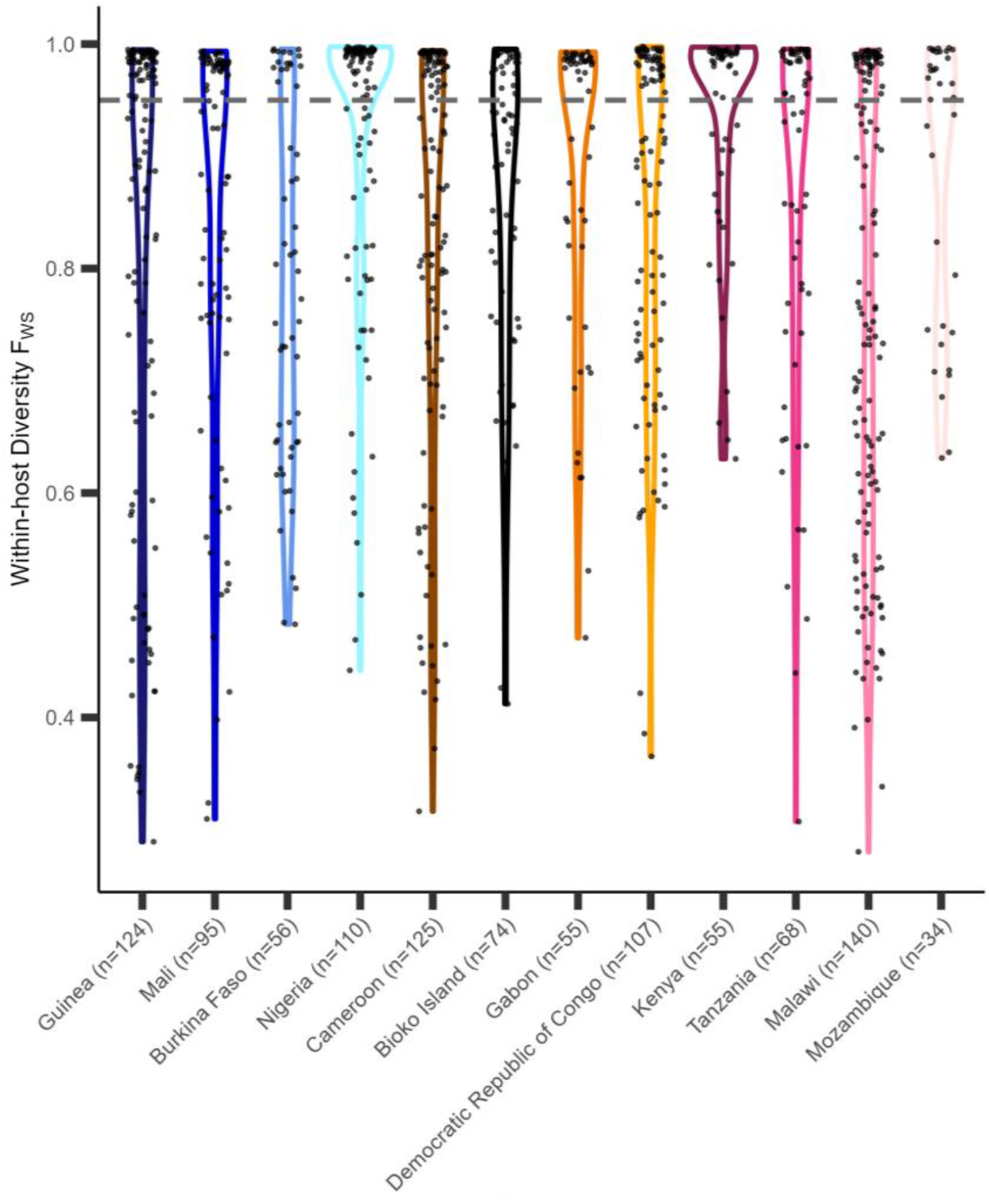
Within-host diversity as measured by *F*_WS_. Y-axis represents *F*_WS_ values of samples between 0 and 1. Isolates with *F*_WS_ value below 0.95 are inferred to be polyclonal (grey dotted line). Color groups denote the African region (West, Central, and East) as assigned by the WHO, and individual colors represent a country were samples were collected (x-axis).

### Bioko Island P. falciparum population is not significantly differentiated from continental neighbors

To determine if Bioko’s island setting has led to an isolated and genetically distinct parasite population relative to its neighboring continental regions, PCA and admixture analysis were conducted. Variance within the sample set could be explained as geographical distance between country groups (West/Central *versus* East = 0.48%; West *versus* Central = 0.24%) (**Figure 4**). Bioko island parasites do not appear to form their own unique population relative to nearby continental parasite populations (*F*_ST Bioko – Cameroon_ = 0.01, p < 0.001; *F*_ST Bioko – Gabon_ = 0.01, p < 0.001). This suggests sufficient connectedness exists with Bioko Island that reduces any isolation effect of a geographical barrier.

**Figure 4.**
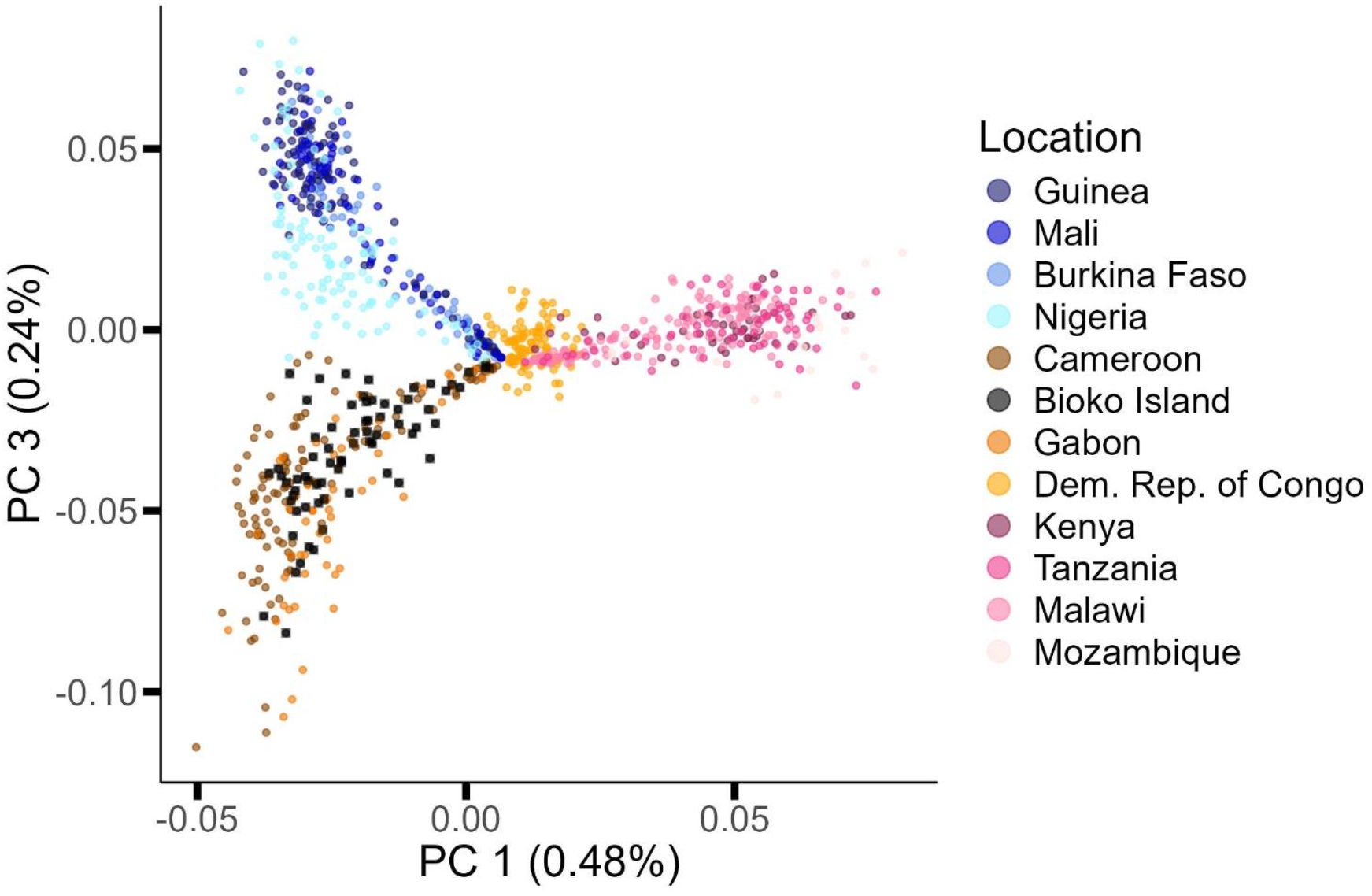
Principal Components Analysis (PCA) of Bioko and African *P. falciparum* strains. PC1 (x-axis) and PC3 (y-axis) illustrate the presence of four main clusters. PC1 separates East Africa from the DRC and both of those regions from countries in both West Africa and the Atlantic coast of Central Africa. In turn, PC3 separates West Africa from Central African populations from on Gulf of Guinea. The DRC sample set is centrally located between all clusters, but does not cluster with other Central African parasites. PC2 is not plotted since clustering pattern is not associated with geography but, instead, reflects the degree of genotype missingness. Color groups denote the African region (West, Central, and East) as assigned by WHO, and colors represent country were samples were collected.

The differentiation seen in the PCA analysis is corroborated by inferred ancestry. An admixture analysis was conducted, which resulted in the inference of the presence of five ancestral populations among the sub-Saharan sample set, with K=5 corresponding to the lowest cross-validation error (**Figure 5**) (Supplementary Figure S3). Bioko Island parasites are characterized by a distinct ancestral population genomic signature that is common to Central-West African: most of the genome represents an ancestral population typical of the Gulf of Guinea (**Figure 5**, green population), followed by representation of ancestral populations more common in West Africa (in orange and yellow in **Figure 5**) and a small genomic proportion typical of Southeast Africa (**Figure 5**, blue and grey). When comparing the ancestral composition between Bioko Island, Cameroon and Gabon, geographical neighbors, the proportion of the blue subpopulation differed (Bioko Island – Cameroon – Gabon: p < 0.001). This subpopulation is predominant in Southeast Africa, and appears to contribute similar levels of ancestry to Bioko and Gabon populations, but not Cameroon (Bioko Island – Gabon: p = 0.43; Bioko Island – Cameroon: p < 0.001; Gabon – Cameroon: p = 0.004). Overall, Bioko parasites appear admixed, with very few (to no) genomes representing a single ancestral source.

**Figure 5.**
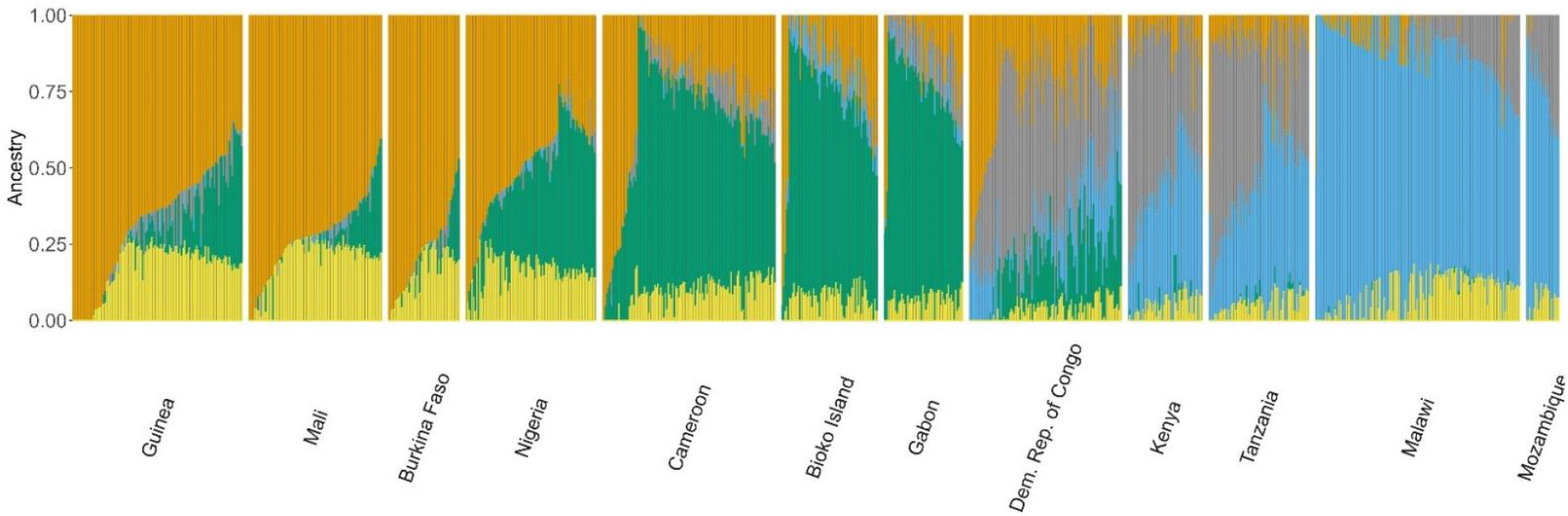
Admixture of Bioko and sub-Saharan parasites. Colors denote each ancestral subpopulation identified for K=5, when cross validation error was lowest. Samples are represented as columns, Y-axis refers to the proportion of ancestry attributable to ancestral subpopulation, in each sample. Orange represents the subpopulation predominant in West Africa, green for Central Africa, grey for Democratic Republic of Congo and East Africa and blue for Southeast Africa. Yellow represents a subpopulation not predominant in any major African region.

### Malabo city parasites differ significantly from those in Baney and Luba in loci associated with cellular adhesion, invasion and sequestration

The human demography of Bioko Island, where the majority of residents live in Malabo city and the surrounding communities are mostly rural, creates the opportunity for a structured *P. falciparum* population on the island. Previous MIS results reported differential measurements of malaria prevalence between urban and rural communities, with higher prevalence in rural communities (8, 19). To determine if multiple parasite populations existed on Bioko Island, we investigated possible genetic differentiation between Bioko-specific geographical subgroups using *F*_ST_. When stratifying samples according to district (Malabo, Baney and Luba), there was some suggestion of overall genetic differentiation between districts, however results were not statistically supported (*F*_ST Malabo vs Baney_ = 0.04, p-value = 0.73; *F*_ST Malabo city vs Luba_ = 0.04, p-value = 0.63). To identify genomic loci contributing to differentiation within Bioko populations, *F*_ST_ per site was calculated between Malabo city, Baney and Luba. Among a total of 163,291 SNPs (Supplementary Figure S4), Baney and Luba consistently differed after 5,000 permutations (p < 0.05) from Malabo city isolates at loci associated with red blood cell invasion (*clag3*.*2* - PF3D7_0302200; *msp* - PF3D7_1035600), hepatocyte invasion (DBL-containing protein - PF3D7_0113800), and parasite sequestration (*var2csa* - PF3D7_1200600).

### Greatest relatedness exists between Bioko and Gabonese parasite populations

To assess the impact of importation on the Bioko *P. falciparum* population, Cameroon and Gabon samples were used as a representative sample set for the suspected source of imported strains to Bioko due to their geographical proximity and admixture results. All samples were deconvoluted, and the predominant strain within each sample used. After merging and filtering, a total of 344,703 SNPs were used. Within Bioko, some strains were highly related (Supplementary Figure S5), but overall mean relatedness of Bioko Island strains was lower than seen within Cameroon or within Gabon (IBD_Bioko_ = 0.003; IBD_Cameroon_ = 0.005; IBD_Gabon_ = 0.006; p < 0.001) (**Figure 6**). Inter-group comparisons illustrated that relatedness of Bioko parasites was twice as high to Gabonese parasites as it was to those from Cameroon (IBD_Bioko-Gabon_ = 0.002; IBD_Bioko-Cameroon_ = 0.001; p < 0.001), and approximately 40% of Bioko parasites formed subpopulations with Gabonese parasites. Ultimately, the *P. falciparum* population within Bioko Island is genetically diverse and composed in largely unrelated parasites, which have marginally higher relatedness to the Gabonese than to the Cameroonian *P. falciparum* population.

**Figure 6.**
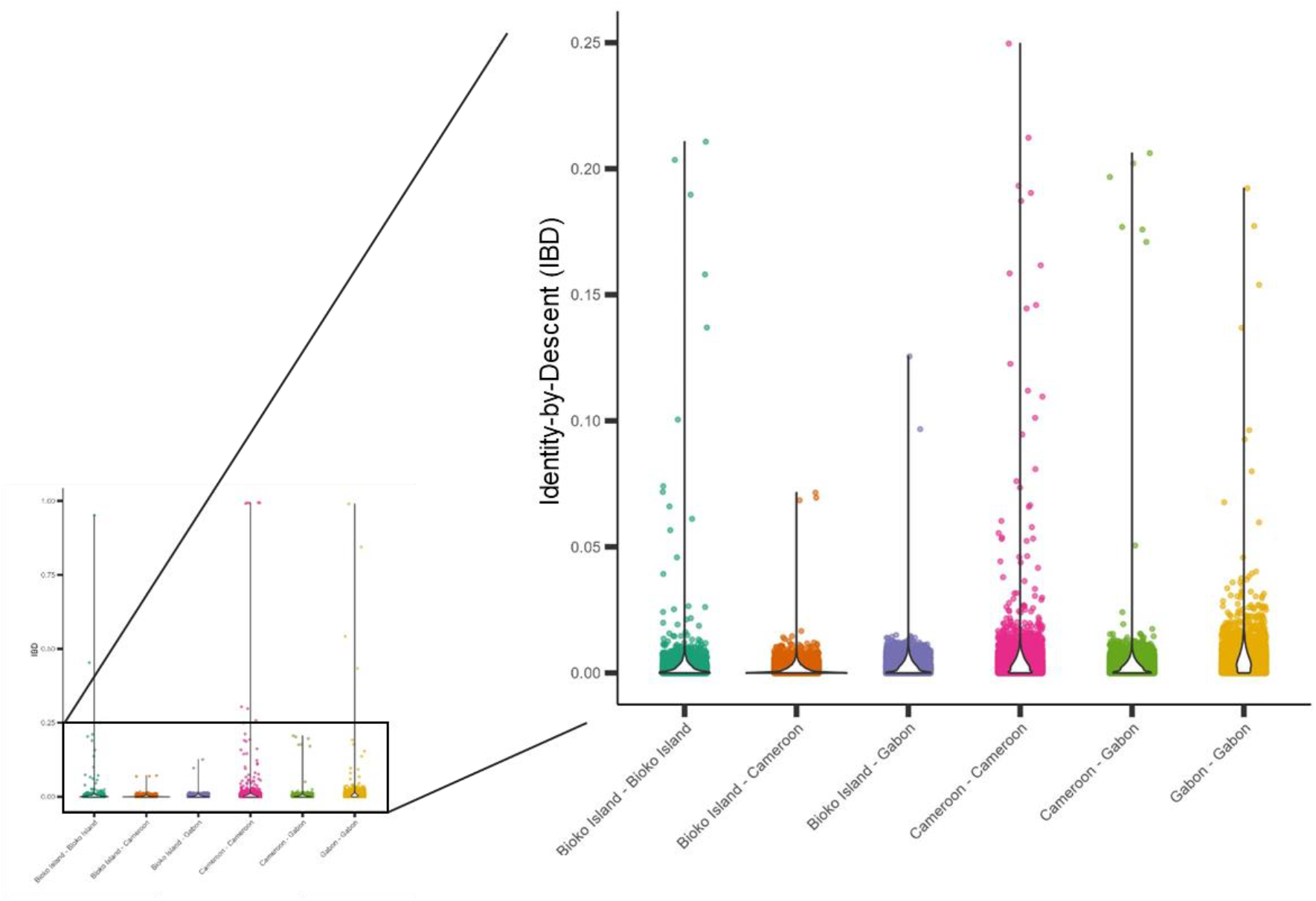
Identity-by-Descent (IBD) of *P. falciparum* pairwise comparisons between Bioko, Cameroon and Gabon. Violin plots illustrate the distribution of IBD values (y-axis) per comparison grouping (x-axis). Most pairwise comparisons are unrelated (IBD < 0.25) and illustrated in the right inset with magnification. All distributions are statistically differentiated from each other (Wilcox rank sum < 0.001).

## Discussion

Considering its central location between Nigeria and the Democratic Republic of Congo (DRC), the two countries globally suffering the highest number of malaria cases in 2022 (27% and 12%, respectively), Bioko island represents an important reference point regarding the disease and potential effect of control interventions within Central Africa (1). As sequencing becomes more accessible, national malaria control programs are incorporating sequencing methodologies as an essential molecular tool to provide improved genotyping accuracy and robustness to supplement traditional epidemiological approaches and inform control and elimination strategies (22, 53-55). As part of the BIMEP’s effort to eliminate malaria from the island of Bioko, WGS data was generated from the 2019 MIS field samples to describe the parasite population at the genomic level.

*P. falciparum* on Bioko Island appears admixed with high genetic diversity and significant connection to the African continent. Malaria genomic epidemiological features were characteristic of endemic areas with moderate transmission (1). For example, most infections were polyclonal; high rates of polyclonality is a proxy measurement of higher transmission intensity since it reflects higher rates of biting by infected mosquitoes (56, 57). Despite large investments in control interventions, complexity of infection on Bioko Island appears similar to other moderate transmission regions in Africa (52, 58, 59). However genetic diversity observations appear similar to high transmission settings of similar contexts, suggesting sustained importation contributes to genetic diversity on Bioko (60, 61). Although human migration likely contributes to on-island transmission and local genetic diversity of *P. falciparum* on Bioko, the observation of many polyclonal infections suggest other factors such as high mosquito biting rates and breeding sites continue to sustain local transmission (62, 63).

The presence, in the genome composition of Bioko samples – as well as in those from Gabon and DRC, of a contribution from an inferred ancestral subpopulation predominant in Southeast Africa hints at a closer connection between those regions that does not extend to Cameroon. This is further supported by the IBD analysis, where Bioko parasites had higher relatedness with Gabonese parasites than with Cameroon. This Gabonese sample set may also be representative of Eastern Rio Muni, a frequent travel destination of off-island travelers (12), as the Gabonese samples used were collected in Wouleu-Ntem province, which has an extensive border with Eastern Equatorial Guinea. There are close cultural and ethnic ties between Equatorial Guinea and Gabon, as the predominant demographic in both countries is the Fang tribe (64-66). Although speculative, this cultural connection may result in increased border crossings and trade in Eastern Rio Muni that then contributes to Bioko’s parasite population when infected travelers return to the island.

On Bioko, there was no apparent population structure and overall relatedness was very low among all pairwise comparisons, suggesting local transmission remains high enough to prevent the detection of clonal expansions or any significant population structure. Although differential SNPs were observed between Bioko district subsets, this is may be due to chance as these occurred on highly polymorphic genomic loci associated with erythrocyte invasion and sequestration (67, 68). Future studies of Bioko Island, and other island-contexts, may benefit from targeted sampling of the most isolated communities to more clearly determine population structure and identify likely local strains. Bioko’s results illustrate the need for extended coverage of malaria control interventions into Rio Muni, and in a broader context, collaboration between malaria endemic countries.

This study had some limitations. No DBS samples have been collected from the continental region of Equatorial Guinea, the most likely source of importation to Bioko, and which may help better distinguish between local and imported strains. Unfortunately, this was unavoidable as, to our knowledge, no MIS has occurred in the Rio Muni since 2011. Also, parasite WGS data was only generated from individuals with positive RDT and fever, to optimize the chances of obtaining good quality sequencing data; however, this bias may have resulted in an over-sampling of complex infections. Sample size was also limited due to resource constraints and may affect results, but is considered minimal since whole genome data provide thousands of data points for analysis. Ultimately, this study follows accepted methodologies (52, 58, 60, 69), and accounts for the limitations within its analysis and interpretations.

Using sWGA, high-quality WGS *P. falciparum* data was generated from DBS collected during the 2019 MIS on Bioko Island, Equatorial Guinea to describe local transmission dynamics and estimate the extent of the local parasite population differentiation from nearby continental regions.

Locally, there did not appear to be any population structure at the genome level. We found no significant genome-wide genetic differentiation between the Bioko Island parasite population and those neighboring countries in mainland Central-West Africa region. Most notably, the flow of importation appeared higher between Bioko and Gabonese parasites, compared to Cameroon, possibly reflecting more frequent human mobility between Bioko and the Eastern Rio Muni. Considerable reductions in island-wide malaria transmission are needed, and importation to the island needs to be addressed to ensure continued progress. Additional sequencing data from parasites on Bioko and the Equatorial Guinea mainland, ideally from both symptomatic and asymptomatic infections, might provide further insights into local transmission dynamics and temporal changes. The current WGS data set provides an informative snapshot of regional relationships among *P. falciparum* populations, successfully sequencing parasites from field samples, and illustrated the effect imported strains may be having on local malaria elimination efforts. Ultimately, for there to be sustained, prolonged progress towards malaria elimination, the BIMEP should consider targeted screening and control interventions for travelers to the island, and if possible, reestablish malaria control in Rio Muni.

## List of abbreviations

BIMEP: Bioko Island Malaria Elimination Project
COI: complexity of infection
DBS: dried blood spot
IBD: identity-by-descent
LD: linkage disequilibrium
MIS: Malaria Indicator Survey
PCA: principal component analysis
PSU: primary sampling units
RBC: red blood cell
SNP: single nucleotide polymorphism
SRA: sequence read archive
sWGA: selective whole genome amplification
WGS: whole genome sequencing

## Declarations

### Ethics approval and consent to participate

Ethics approval for the 2019 MIS was provided by the Equatorial Guinea Ministry of Health and Social Welfare and the ethics committee of the London School of Hygiene and Tropical Medicine (approval number 5556). Written informed consent was sought from each participating adult and on behalf of participating children under 18 years of age.

### Consent for publication

Not applicable

### Availability of data and materials

Sequences generated from this study have been deposited in GenBank under BioProject ID PRJNA1193657. The survey data collected during the MIS for the current study are available from the corresponding author(s) upon reasonable request.

The list of publicly acquired sequences can be found in Supplementary File 2.

### Competing Interests

The authors declare that they have no competing interests.

### Funding

Work related to the BIMEP activities was funded by a private sector consortium led by Marathon Oil Corporation and the Government of Equatorial Guinea. The funders had no role in study design, data collection, data analysis, data interpretation, the decision to publish, or preparation of the manuscript. Work related to the generation of WGS data was funded by the National Institutes of Health (NIH) awards R01 AI141900 to JCS.

### Author’s contributions

TCS conceived the study, performed all DNA extractions, qPCR assays and sWGA, conducted analyses, interpreted results and wrote the paper. AD supported the study design and methodology, contributed to the interpretation of results and edited the manuscript. BG supported analyses of relatedness and reviewed results and guided interpretations. BS and SJ advised and supported all laboratory procedures and edited the paper. MRR sponsored and oversaw the 2019 MIS. OTD managed the 2019 MIS teams collecting all field samples, organized sample logistics and edited the paper. CAG contributed to the study design, supported interpretation of 2019 MIS epidemiological data, assisted in generating Bioko Island figures, and edited the paper. GAG contributed fundamental funding and resources to the study, contributed to the study design, and edited the paper. CD conceived of the study, contributed essential resources to the study and wrote the paper. JCS conceived of the study, contributed to the study design, interpretation of results, contributed funding and resources and wrote the paper. All authors read and approved the final version of the manuscript.

## Acknowledgements

This work was supported by the National Malaria Control Programme and the Ministry of Health and Social Welfare of Equatorial Guinea, and MCD Global Health through the Bioko Island Malaria Elimination Project (BIMEP). Specifically, we would like to thank the entire BIMEP team for their efforts conducting the 2019 MIS. Ultimately, we would like to thank Marathon Oil, Noble Energy, AMPCO (Atlantic Methanol Production Company), and the Ministry of Mines and Energy of Equatorial Guinea for their continued efforts and support for malaria control on Bioko Island. This work was also supported by the National Institutes of Health through grant R01AI141900. We thank the staff of the IGS’s Maryland Genomics for genomic DNA library construction and sequencing.

## References

1. WHO. World malaria report 2023: World Health Organization; 2023.

2. WHO. WHO certifies Cabo Verde as malaria-free, marking a historic milestone in the fight against malaria. 2024 [Available from: https://www.who.int/news/item/12-01-2024-who-certifies-cabo-verde-as-malaria-free--marking-a-historic-milestone-in-the-fight-against-malaria.

3. Senaratne R, Singh PK. Against the odds, Sri Lanka eliminates malaria. The Lancet. 2016;388(10049):1038–9.

4. Ali MH, Kitau J, Ali AS, Al-Mafazy AW, Tegegne SG, Ussi O, et al. Malaria elimination in Zanzibar: where next? Pan Afr Med J. 2023;45(Suppl 1):7.

5. Wang Y, Li M, Guo W, Deng C, Zou G, Song J. Burden of Malaria in Sao Tome and Principe, 1990–2019: Findings from the Global Burden of Disease Study 2019. International Journal of Environmental Research and Public Health. 2022;19(22):14817.

6. Markwalter CF, Menya D, Wesolowski A, Esimit D, Lokoel G, Kipkoech J, et al. Plasmodium falciparum importation does not sustain malaria transmission in a semi-arid region of Kenya. PLOS Global Public Health. 2022;2(8):e0000807.

7. Wesolowski A, Eagle N, Tatem AJ, Smith DL, Noor AM, Snow RW, et al. Quantifying the impact of human mobility on malaria. Science. 2012;338(6104):267–70.

8. Cook J, Hergott D, Phiri W, Rivas MR, Bradley J, Segura L, et al. Trends in parasite prevalence following 13 years of malaria interventions on Bioko island, Equatorial Guinea: 2004–2016. Malaria Journal. 2018;17(1):62.

9. García GA, Janko M, Hergott DEB, Donfack OT, Smith JM, Mba Eyono JN, et al. Identifying individual, household and environmental risk factors for malaria infection on Bioko Island to inform interventions. Malaria Journal. 2023;22(1):72.

10. Nchama VUNN, Said AH, Mtoro A, Bidjimi GO, Owono MA, Maye ERM, et al. Incidence of Plasmodium falciparum malaria infection in 6-month to 45-year-olds on selected areas of Bioko Island, Equatorial Guinea. Malaria Journal. 2021;20(1):322.

11. Bradley J, Monti F, Rehman AM, Schwabe C, Vargas D, Garcia G, et al. Infection importation: a key challenge to malaria elimination on Bioko Island, Equatorial Guinea. Malaria Journal. 2015;14(1):46.

12. Guerra CA, Kang SY, Citron DT, Hergott DEB, Perry M, Smith J, et al. Human mobility patterns and malaria importation on Bioko Island. Nature Communications. 2019;10(1):2332.

13. Citron DT, Guerra CA, García GA, Wu SL, Battle KE, Gibson HS, et al. Quantifying malaria acquired during travel and its role in malaria elimination on Bioko Island. Malaria Journal. 2021;20(1):359.

14. Hergott DEB, Guerra CA, García GA, Mba Eyono JN, Donfack OT, Iyanga MM, et al. Impact of six-month COVID-19 travel moratorium on Plasmodium falciparum prevalence on Bioko Island, Equatorial Guinea. Nature Communications. 2024;15(1):8285.

15. Rehman AM, Mann AG, Schwabe C, Reddy MR, Roncon Gomes I, Slotman MA, et al. Five years of malaria control in the continental region, Equatorial Guinea. Malaria Journal. 2013;12(1):154.

16. Atuh NI, Anong DN, Jerome F-C, Oriero E, Mohammed NI, D’Alessandro U, et al. High genetic complexity but low relatedness in Plasmodium falciparum infections from Western Savannah Highlands and coastal equatorial Lowlands of Cameroon. Pathogens and Global Health. 2022;116(7):428–37.

17. Mvé-Ondo B, Nkoghe D, Arnathau C, Rougeron V, Bisvigou U, Mouele LY, et al. Genetic diversity of Plasmodium falciparum isolates from Baka Pygmies and their Bantu neighbours in the north of Gabon. Malaria Journal. 2015;14(1):395.

18. Fries B, Guerra CA, García GA, Wu SL, Smith JM, Oyono JNM, et al. Measuring the accuracy of gridded human population density surfaces: A case study in Bioko Island, Equatorial Guinea. PLOS ONE. 2021;16(9):e0248646.

19. Guerra CA, Citron DT, García GA, Smith DL. Characterising malaria connectivity using malaria indicator survey data. Malaria Journal. 2019;18(1):440.

20. Neafsey DE, Taylor AR, MacInnis BL. Advances and opportunities in malaria population genomics. Nature Reviews Genetics. 2021;22(8):502–17.

21. Holzschuh A, Lerch A, Gerlovina I, Fakih BS, Al-mafazy A-wH, Reaves EJ, et al. Multiplexed ddPCR-amplicon sequencing reveals isolated Plasmodium falciparum populations amenable to local elimination in Zanzibar, Tanzania. Nature Communications. 2023;14(1):3699.

22. WHO. Global technical strategy for malaria 2016–2030, 2021 update. J Geneva: World Health Organization. 2021:1–40.

23. Amambua-Ngwa A, Amenga-Etego L, Kamau E, Amato R, Ghansah A, Golassa L, et al. Major subpopulations of Plasmodium falciparum in sub-Saharan Africa. 2019;365(6455):813–6.

24. Oberstaller J, Zoungrana L, Bannerman CD, Jahangiri S, Dwivedi A, Silva JC, et al. Integration of population and functional genomics to understand mechanisms of artemisinin resistance in Plasmodium falciparum. International Journal for Parasitology: Drugs and Drug Resistance. 2021;16:119–28.

25. Rocamora F, Winzeler EA. Genomic Approaches to Drug Resistance in Malaria. Annual Review of Microbiology. 2020;74(Volume 74, 2020):761–86.

26. Mensah BA, Akyea-Bobi NE, Ghansah A. Genomic approaches for monitoring transmission dynamics of malaria: A case for malaria molecular surveillance in Sub–Saharan Africa. Frontiers in Epidemiology. 2022;2.

27. García GA, Hergott DEB, Phiri WP, Perry M, Smith J, Osa Nfumu JO, et al. Mapping and enumerating houses and households to support malaria control interventions on Bioko Island. Malaria Journal. 2019;18(1):283.

28. MalariaGEN, Abdel Hamid M, Abdelraheem M, Acheampong D, Ahouidi A, Ali M, et al. Pf7: an open dataset of Plasmodium falciparum genome variation in 20,000 worldwide samples [version 1; peer review: 3 approved]. Wellcome Open Research. 2023;8(22).

29. Zainabadi K, Adams M, Han ZY, Lwin HW, Han KT, Ouattara A, et al. A novel method for extracting nucleic acids from dried blood spots for ultrasensitive detection of low-density Plasmodium falciparum and Plasmodium vivax infections. Malaria Journal. 2017;16(1):377.

30. Shah Z, Adams M, Moser KA, Shrestha B, Stucke EM, Laufer MK, et al. Optimization of parasite DNA enrichment approaches to generate whole genome sequencing data for Plasmodium falciparum from low parasitaemia samples. Malaria Journal. 2020;19:1–10.

31. Oyola SO, Ariani CV, Hamilton WL, Kekre M, Amenga-Etego LN, Ghansah A, et al. Whole genome sequencing of Plasmodium falciparum from dried blood spots using selective whole genome amplification. Malaria Journal. 2016;15(1):597.

32. Moser KA, Drábek EF, Dwivedi A, Stucke EM, Crabtree J, Dara A, et al. Strains used in whole organism Plasmodium falciparum vaccine trials differ in genome structure, sequence, and immunogenic potential. Genome Medicine. 2020;12(1):6.

33. Langmead B, Salzberg SL. Fast gapped-read alignment with Bowtie 2. J Nature Methods. 2012;9(4):357.

34. DePristo MA, Banks E, Poplin R, Garimella KV, Maguire JR, Hartl C, et al. A framework for variation discovery and genotyping using next-generation DNA sequencing data. J Nature genetics. 2011;43(5):491.

35. Van der Auwera GA, Carneiro MO, Hartl C, Poplin R, del Angel G, Levy-Moonshine A, et al. From FastQ Data to High-Confidence Variant Calls: The Genome Analysis Toolkit Best Practices Pipeline. Curr Protoc Bioinform. 2013;43(1):11.0.1-.0.33.

36. Quinlan AR, Hall IM. BEDTools: a flexible suite of utilities for comparing genomic features. Bioinformatics. 2010;26(6):841–2.

37. Karamoko Niaré BG, Jeffrey A Bailey. An Optimized GATK4 Pipeline for Plasmodium falciparum Whole Genome Sequencing Variant Calling and Analysis. Malaria Journal. 2023;PREPRINT.

38. Manske M, Miotto O, Campino S, Auburn S, Almagro-Garcia J, Maslen G, et al. Analysis of Plasmodium falciparum diversity in natural infections by deep sequencing. Nature. 2012;487(7407):375–9.

39. Zheng X, Levine D, Shen J, Gogarten SM, Laurie C, Weir BS. A high-performance computing toolset for relatedness and principal component analysis of SNP data. Bioinformatics. 2012;28(24):3326–8.

40. Zheng X, Gogarten SM, Lawrence M, Stilp A, Conomos MP, Weir BS, et al. SeqArray—a storage-efficient high-performance data format for WGS variant calls. Bioinformatics. 2017;33(15):2251–7.

41. Weir BS, Cockerham CC. Estimating F-Statistics for the Analysis of Population Structure. Evolution. 1984;38(6):1358–70.

42. Schaffner SF, Taylor AR, Wong W, Wirth DF, Neafsey DE. hmmIBD: software to infer pairwise identity by descent between haploid genotypes. Malaria Journal. 2018;17(1):196.

43. Zhu SJ, Almagro-Garcia J, McVean G. Deconvolution of multiple infections in Plasmodium falciparum from high throughput sequencing data. Bioinformatics. 2017;34(1):9–15.

44. Rosvall M, Bergstrom CT. Maps of random walks on complex networks reveal community structure. Proceedings of the National Academy of Sciences. 2008;105(4):1118–23.

45. Guo B, Borda V, Laboulaye R, Spring MD, Wojnarski M, Vesely BA, et al. Strong positive selection biases identity-by-descent-based inferences of recent demography and population structure in Plasmodium falciparum. Nature Communications. 2024;15(1):2499.

46. Shetty AC, Jacob CG, Huang F, Li Y, Agrawal S, Saunders DL, et al. Genomic structure and diversity of Plasmodium falciparum in Southeast Asia reveal recent parasite migration patterns. Nature Communications. 2019;10(1):2665.

47. Pebesma E. Simple Features for R: Standardized Support for Spatial Vector Data. The R Journal. 2018;10(1):439–46.

48. Miles A, Iqbal Z, Vauterin P, Pearson R, Campino S, Theron M, et al. Indels, structural variation, and recombination drive genomic diversity in Plasmodium falciparum. Genome Res. 2016;26(9):1288–99.

49. Otto TD, Böhme U, Sanders M, Reid A, Bruske EI, Duffy CW, et al. Long read assemblies of geographically dispersed Plasmodium falciparum isolates reveal highly structured subtelomeres. Wellcome Open Res. 2018;3:52.

50. Gray K-A, Dowd S, Bain L, Bobogare A, Wini L, Shanks GD, et al. Population genetics of Plasmodium falciparum and Plasmodium vivax and asymptomatic malaria in Temotu Province, Solomon Islands. Malaria Journal. 2013;12(1):429.

51. Arez AP, Snounou G, Pinto J, Sousa CA, Modiano D, Ribeiro H, et al. A clonal Plasmodium falciparum population in an isolated outbreak of malaria in the Republic of Cabo Verde. Parasitology. 1999;118(4):347–55.

52. Moss S, Mańko E, Vasileva H, Da Silva ET, Goncalves A, Osborne A, et al. Population dynamics and drug resistance mutations in Plasmodium falciparum on the Bijagós Archipelago, Guinea-Bissau. Scientific Reports. 2023;13(1):6311.

53. Dalmat R, Naughton B, Kwan-Gett TS, Slyker J, Stuckey EM. Use cases for genetic epidemiology in malaria elimination. Malaria Journal. 2019;18(1):163.

54. Golumbeanu M, Edi CAV, Hetzel MW, Koepfli C, Nsanzabana C. Bridging the Gap from Molecular Surveillance to Programmatic Decisions for Malaria Control and Elimination. The American Journal of Tropical Medicine and Hygiene. 2023:tpmd220749.

55. Tessema SK, Raman J, Duffy CW, Ishengoma DS, Amambua-Ngwa A, Greenhouse B. Applying next-generation sequencing to track falciparum malaria in sub-Saharan Africa. Malaria Journal. 2019;18(1):268.

56. Nkhoma SC, Nair S, Al-Saai S, Ashley E, McGready R, Phyo AP, et al. Population genetic correlates of declining transmission in a human pathogen. Molecular Ecology. 2013;22(2):273–85.

57. Neafsey DE, Volkman SK. Malaria Genomics in the Era of Eradication. J Cold Spring Harbor Perspectives in Medicine. 2017;7(8).

58. Roh ME, Tessema SK, Murphy M, Nhlabathi N, Mkhonta N, Vilakati S, et al. High Genetic Diversity of Plasmodium falciparum in the Low-Transmission Setting of the Kingdom of Eswatini. J Infect Dis. 2019;220(8):1346–54.

59. Amambua-Ngwa A, Jeffries D, Mwesigwa J, Seedy-Jawara A, Okebe J, Achan J, et al. Long-distance transmission patterns modelled from SNP barcodes of Plasmodium falciparum infections in The Gambia. Scientific Reports. 2019;9(1):13515.

60. Kassegne K, Komi Koukoura K, Shen H-M, Chen S-B, Fu H-T, Chen Y-Q, et al. Genome-Wide Analysis of the Malaria Parasite Plasmodium falciparum Isolates From Togo Reveals Selective Signals in Immune Selection-Related Antigen Genes. Frontiers in Immunology. 2020;11.

61. Papa Mze N, Bogreau H, Diedhiou CK, Herdell V, Rahamatou S, Bei AK, et al. Genetic diversity of Plasmodium falciparum in Grande Comore Island. Malaria Journal. 2020;19(1):320.

62. Guerra CA, Fuseini G, Donfack OT, Smith JM, Ondo Mifumu TA, Akadiri G, et al. Malaria outbreak in Riaba district, Bioko Island: lessons learned. Malaria Journal. 2020;19(1):277.

63. Reddy MR, Overgaard HJ, Abaga S, Reddy VP, Caccone A, Kiszewski AE, et al. Outdoor host seeking behaviour of Anopheles gambiae mosquitoes following initiation of malaria vector control on Bioko Island, Equatorial Guinea. Malaria Journal. 2011;10(1):184.

64. Stokes J. Encyclopedia of the Peoples of Africa and the Middle East: Infobase Publishing; 2009.

65. CIA. Equatorial Guinea: People and Society 2024 [Available from: https://www.cia.gov/the-world-factbook/countries/equatorial-guinea/.

66. CIA. Cameroon: People and Society 2024 [Available from: https://www.cia.gov/the-world-factbook/countries/cameroon/.

67. Mobegi VA, Duffy CW, Amambua-Ngwa A, Loua KM, Laman E, Nwakanma DC, et al. Genome-wide analysis of selection on the malaria parasite Plasmodium falciparum in West African populations of differing infection endemicity. Mol Biol Evol. 2014;31(6):1490–9.

68. Tomlinson A, Semblat J-P, Gamain B, Chêne A. VAR2CSA-Mediated Host Defense Evasion of Plasmodium falciparum Infected Erythrocytes in Placental Malaria. Frontiers in Immunology. 2021;11.

69. Vanheer LN, Mahamar A, Manko E, Niambele SM, Sanogo K, Youssouf A, et al. Genome-wide genetic variation and molecular surveillance of drug resistance in Plasmodium falciparum isolates from asymptomatic individuals in Ouélessébougou, Mali. Scientific Reports. 2023;13(1):9522.

